# Temporal Integration of Weak Acoustic Stress Leads to Delayed Cellular Responses

**DOI:** 10.64898/2026.10.18.759952

**Authors:** Young Hyun Noh, I-Ting Ho, Melody Kosters, Keith Runge, Xiaoyun Ding, Pierre A. Deymier, Minkyu Kim

## Abstract

Sound is a pervasive physical stimulus with beneficial or adverse effects depending on exposure. While controlled sound-based interventions such as ultrasound have demonstrated clinical utility, epidemiological studies link chronic environmental noise to increased cardiovascular and metabolic risk. Despite these associations, how weak sound is sensed and integrated at the cellular level remains unclear. Here, we investigate cellular responsiveness to acoustic pulse-wave stimulation in human endothelial cells, using intracellular calcium changes as a readout of cellular activation. Cellular responses to acoustic stimulation were compared with over-threshold mechanical stimulation. Over-threshold mechanical stimulation induced immediate responses, whereas single sub-threshold acoustic exposures produced no detectable response. In contrast, sustained sub-threshold acoustic stimulation over minutes produced delayed activation, with onset times dependent on stimulus intensity. These results demonstrate that cells integrate weak physical inputs over time, revealing a delayed activation regime under sustained sub-threshold exposure.

## 1. Introduction

Sound is a ubiquitous physical stimulus in daily life, with biological effects that depend on exposure intensity, frequency, and duration. Increasing evidence supports a dual role for sound: controlled sound-based interventions, including ultrasound, can provide therapeutic and diagnostic benefits (*1-4*), whereas unintended and prolonged exposure to environmental noise is statistically associated with elevated cardiovascular and metabolic risks (*5, 6*). Despite these systemic observations, how sound is sensed, processed, and translated into biological responses at the cellular level remains incompletely elucidated.

Recent studies suggest that sound can act as a physical stimulus to cells even within the audible frequency range (*7-9*). Reported outcomes include changes in gene expression and mechanosensory signaling pathways, indicating that cells can perceive acoustic or vibrational cues from their environment (*8, 9*). Yet, these observations largely reflect downstream endpoints and do not resolve the early intracellular signaling events that couple external acoustic stimuli to cellular responses. Accordingly, understanding how physical stimuli are initially sensed and transduced at the cellular level requires examining early intracellular signaling processes, including calcium signaling (*10, 11*). Calcium signaling is a sensitive readout of early cellular responses and a central regulator of mechanotransduction, intercellular communication, and signaling pathways that operate both upstream and downstream to modulate cellular adaptation and function (*12, 13*). Calcium dynamics have been extensively characterized under strong mechanical and high-dose chemical perturbations (*14-16*). However, it remains unclear how weak acoustic stress influences calcium signaling during prolonged exposure, a regime potentially relevant to exposure to long-term environmental noise.

Weak or sub-threshold physical stimuli are often assumed to have negligible biological impact because they do not elicit readily detectable cellular responses under typical experimental conditions. However, the absence of an acute, readily measurable response does not preclude the possibility that cells integrate repeated sub-threshold inputs over time and eventually cross a functional activation threshold (*17, 18*). This raises the question of whether prolonged exposure to low-intensity acoustic stress, even when individual stimuli are insufficient to elicit a prompt response, can nonetheless trigger measurable cellular responses through temporal integration. Calcium signaling provides a sensitive readout to test this hypothesis by capturing delayed or cumulative intracellular responses that may be missed in short-term assays. Conceptually, this possibility aligns with stress-integration paradigms, in which sustained perturbations can be converted into signaling outputs (*19, 20*).

Herein, we investigated how repeated sub-threshold acoustic pulse-wave stimulation influences calcium signaling in human endothelial cells. We compared the temporal dynamics of intracellular calcium signaling across distinct modalities, using over-threshold mechanical stimulation as a positive control and sub-threshold light stimulation as an orthogonal perturbation. By focusing on early calcium signaling dynamics, this study establishes a framework for understanding how prolonged acoustic exposure may be encoded at the cellular level and propagated to broader physiological responses over time.

## 2. Results

### 2.1. Experimental platform for controlled physical stimulation and calcium imaging

To investigate cellular responses to weak and sustained physical stimuli, we established an experimental platform that enabled controlled delivery of surface-propagating pulse-waves while remaining compatible with live-cell calcium imaging (**Fig. 1**). This platform allowed direct comparison of intracellular calcium dynamics across distinct physical stimulation modalities under consistent imaging and cell-culture conditions. Cells were cultured in a customized PDMS chamber attached to a glass coverslip, which served as the transparent substrate for live-cell fluorescence imaging and defined the cell-culture reservoir. The same PDMS structure incorporated a dedicated mount that fixed the position and distance of a piezoelectric transducer relative to the cell-culture region. Pulse-waves generated at this location propagated laterally along the coverslip, allowing cells to experience surface-mediated physical stimulation without direct contact with the transducer. Within this geometry, stimulation parameters—including pulse repetition frequency, pulse duty cycle defining the temporal on–off structure, and amplitude controlled via the transducer input voltage—were independently modulated. This enabled control over the pulse-wave profile and the strength of the applied stimulation. Acoustic absorbers were incorporated upstream and downstream of the cell-culture region to suppress unintended wave reflections and back-propagation from external boundaries. These absorbers also decoupled the coverslip from external mechanical vibrations, enabling a mechanically isolated environment that preserved the applied pulse-wave profile. Surface-wave propagation and temporal characteristics at the cell-culture region were independently validated using laser Doppler vibrometry (LDV) in the absence of cells, confirming consistent delivery of the applied pulse-wave stimuli (**Fig. S1**). By maintaining consistent cell-culture geometry and imaging conditions, this platform enabled direct comparison of the temporal dynamics of intracellular calcium signaling across selected stimulation modalities.

**Fig. 1.**
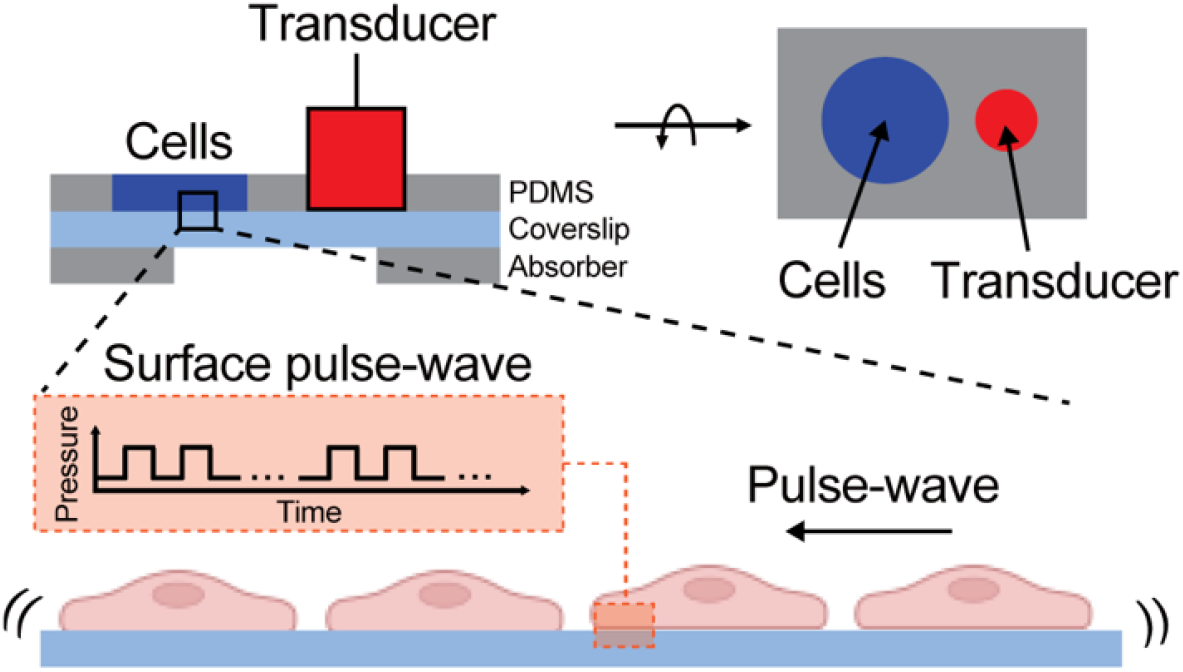
Schematic of surface-mediated pulse-wave stimulation of human endothelial cells. A piezoelectric transducer positioned above a glass coverslip generates pulse-waves that propagate laterally along the coverslip toward a spatially separated cell-culture region. The cross-sectional view (top left) shows the layered configuration of the transducer, coverslip, PDMS block, and acoustic absorber, which together define the stimulation geometry and minimize wave reflections. The top-down view (top right) shows the lateral separation between the transducer and the cell-culture region. The temporal structure of the applied pulse-wave is illustrated schematically (middle) and independently validated by laser Doppler vibrometry (Fig. S1). The bottom panel depicts surface-wave propagation along the substrate beneath adherent cells.

### 2.2. Delayed calcium signaling driven by temporal integration of weak physical stimulation

To establish a reference for calcium responses under clearly over-threshold conditions, we first examined intracellular calcium dynamics in human umbilical vein endothelial cells subjected to direct mechanical stimulation using a microneedle. Experiments were performed using the same imaging platform and culture geometry shown in **Fig. 1**, with the piezoelectric transducer omitted and mechanical perturbation applied locally to individual cells. Fluorescence imaging was conducted under low-intensity conditions (488 nm, 0.15 mW·cm^−2^), which did not elicit detectable calcium activation over the duration of the experiment in the absence of stimulation (**Fig. 2A**). Calcium signaling was monitored as increases in intracellular fluorescence, reflecting rapid calcium release from endoplasmic reticulum stores upon direct mechanical perturbation. Upon microneedle contact, a rapid increase in intracellular calcium was observed in the directly stimulated cell, with detectable activation occurring within ∼0.1 s of stimulation onset (**Fig. 2B**). In addition to the directly perturbed cell, calcium activation propagated to neighboring cells within the endothelial monolayer (**Fig. S2**). These responses are consistent with previously reported endothelial calcium signaling induced by direct mechanical perturbation (*14, 15*) and confirm that the experimental platform reliably captures rapid cellular activation induced by over-threshold stimulation.

**Fig. 2.**
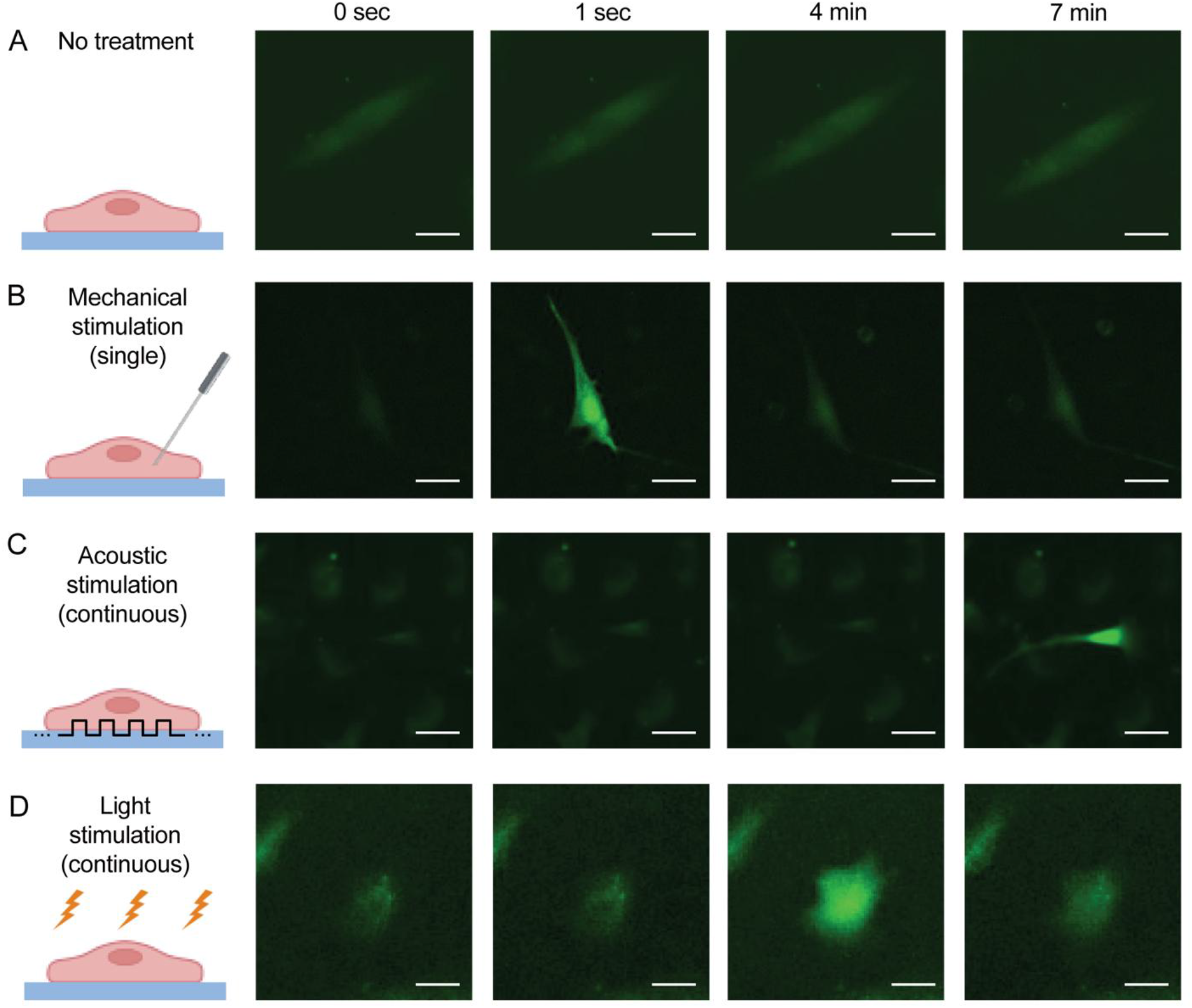
Intracellular calcium dynamics in endothelial cells under distinct physical stimulation modalities. Representative time-lapse fluorescence images of intracellular calcium signaling in human umbilical vein endothelial cells (HUVECs) loaded with a calcium-sensitive indicator, Calbryte 520AM. Images are shown at the indicated time points (0 s, 1 s, 4 min, and 7 min) following stimulus onset. (**A**) Unstimulated control cells showing basal fluorescence over time. (**B**) Direct mechanical stimulation applied by a single microneedle contact, producing a rapid increase in intracellular calcium detectable within 1 s in the stimulated cell. (**C**) Continuous sub-threshold acoustic stimulation delivered as surface-propagating pulse-waves (10 kHz, 170 V input, 70% duty cycle), with intracellular calcium activation at later time points. (**D**) Continuous sub-threshold light stimulation (488 nm, 1.0 mW·cm−²), with calcium activation at later time points. Cropped regions shown here were extracted from larger fields of view; corresponding full-field images are provided in Fig. S3. Scale bar, 20 µm.

We next examined intracellular calcium responses under sub-threshold acoustic stimulation delivered as surface-propagating pulse-waves that impose weak mechanical perturbations at the cell–substrate interface. Pulse-wave stimulation was applied using a pulse repetition frequency of 10 kHz with a 70% duty cycle at 170 V, selected to impose weak mechanical input at the cell-culture region. Attenuation through the coverslip, buffer, and intervening PDMS structure substantially reduced the surface-wave amplitude relative to the transducer output (**Fig. 1**), while preserving directional surface-wave propagation, as confirmed by LDV measurements (**Fig. S1**). Under these conditions, no detectable increase in intracellular calcium was observed within the first minute of stimulation across the observed cell population (**Fig. 2C**). This lack of a rapid response was consistently observed across independent experiments, indicating that weak surface-mediated mechanical stimulation at sub-threshold intensity does not elicit detectable calcium activation on short timescales.

We then investigated whether sustained exposure to sub-threshold pulse-wave stimulation could elicit intracellular calcium responses over longer timescales. With continued stimulation, delayed calcium activation emerged after several minutes within the observed cell population (**Fig. 2C**). Activation time was defined as the time to the maximum intracellular calcium signal observed in the fluorescence trace of an individual cell. These responses were reproducible, although the specific responding cells and their spatial locations varied across imaging fields, with first-cell activation observed at *t* = 403.3 ± 25.2 s (*n* = 3). In contrast to over-threshold mechanical stimulation, the delayed calcium responses remained confined to individual cells and did not propagate to neighboring cells. Together, these observations show that sub-threshold acoustic surface-mediated stimulation does not induce rapid calcium signaling, yet sustained exposure can give rise to delayed intracellular calcium activation.

To assess whether delayed calcium activation was specific to surface-mediated physical inputs or extended beyond acoustically generated stimulation, we additionally examined intracellular calcium dynamics under blue light stimulation as an orthogonal physical perturbation. Blue light was used to match the fluorescence excitation wavelength of the calcium indicator used for imaging. The light intensity (488 nm, 1.0 mW·cm^−2^) was set such that no detectable calcium activation occurred on short timescales following stimulus onset (**Fig. 2D**). Similar to sub-threshold pulse-wave stimulation, sustained illumination induced calcium activation after approximately one minute within the observed cell population, exhibiting delayed onset behavior. The response timing was reproducible across independent experiments (*t* = 244.2 ± 54.1 s; *n* = 3), although the responding cells varied across imaging fields. Light-induced calcium activation also remained confined to individual cells and did not propagate to neighboring cells. Altogether, these results indicate that delayed calcium activation under sustained sub-threshold exposure is not limited to acoustically generated surface-mediated stimulation, but reflects a more general cellular capacity to integrate weak physical stimuli over time.

### 2.3. Intensity-dependent modulation of delayed calcium signaling

After establishing delayed calcium activation under sustained sub-threshold stimulation, we further examined whether the timing of these responses depended on the amplitude of the applied physical input. For sub-threshold acoustic stimulation, pulse-wave amplitude was systematically varied by adjusting the transducer input voltage (140 V, 170 V, and 200 V), while maintaining stimulation pulse repetition frequency (10 kHz) and duty cycle (70%). Across all tested amplitudes, no immediate calcium activation was observed following stimulation onset. However, the latency to calcium activation varied with applied voltage, with higher pulse-wave amplitudes associated with earlier onset of delayed calcium responses (**Fig. 3A**). Note that fluorescence imaging alone did not lead to calcium activation. Quantitatively, the time to first detectable calcium activation decreased from 803.3 ± 98.7 s at 140 V to 403.3 ± 25.2 s at 170 V, and further to 128.3 ± 65.5 s at 200 V (*n* = 3; **Fig. 3B**). These results show that under sub-threshold acoustic stimulation, the timing of delayed calcium activation depends on pulse-wave amplitude rather than exhibiting a fixed latency.

**Fig. 3.**
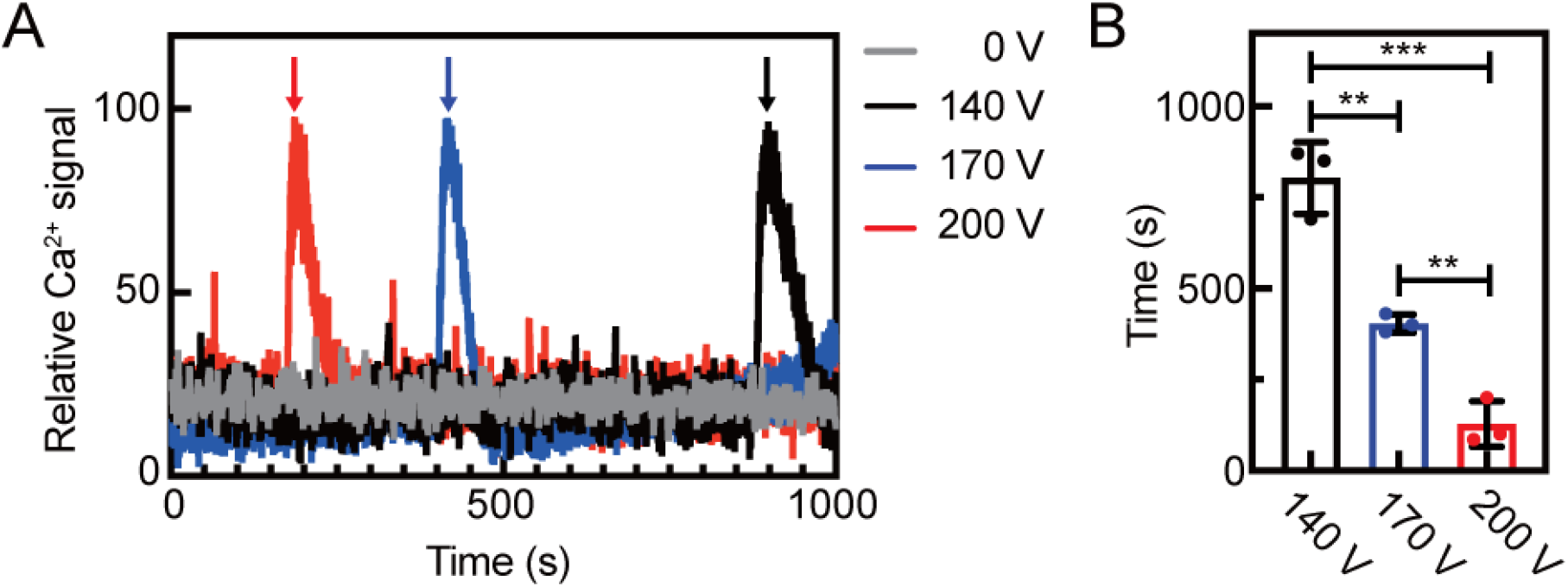
Intensity-dependent delay of calcium activation under sub-threshold acoustic stimulation. (**A**) Representative fluorescence time traces of intracellular calcium responses in HUVECs subjected to sustained sub-threshold acoustic stimulation. Traces are shown under three pulse-wave input voltages (0, 140, 170, 200 V; 10 kHz; 70% duty cycle). Fluorescence imaging was performed using 488 nm blue light at an intensity of 0.15 mW·cm^−2^, which did not elicit detectable calcium activation without acoustic stimulation (gray). Relative Ca^2+^ signals were calculated from background-subtracted fluorescence and subsequently min-max normalized, with values scaled 0 and 100, respectively. Calcium activation occurred with elayed onset under all conditions, with earlier onset at higher pulse-wave intensities. (**B**) Quantification of latency to first detectable calcium activation under sub-threshold acoustic stimulation. Higher input voltages significantly reduced activation latency (140 V: 803.3 ± 98.7 s; 170 V: 403.3 ± 25.2 s; 200 V: 128.3 ± 65.5 s; mean ± s.d., *n* = 3). Latency was defined as the time required to reach peak calcium signal (indicated by arrows in panel a). Statistical significance was determined by one-way ANOVA followed by Tukey’s post hoc multiple-comparisons test to account for multiple hypothesis testing. Note: *n* = 3; \*\**p* < 0.01; \*\*\**p* < 0.001.

A similar intensity-dependent effect was observed under sub-threshold light stimulation. When blue light intensity was varied while maintaining constant illumination wavelength and exposure geometry, calcium activation remained delayed but occurred progressively earlier with increasing light intensity (**Fig. 4A**). Compared to acoustic stimulation, calcium elevation under light stimulation spanned a broader temporal range. No calcium activation was observed at 0.15 mW·cm^-2^. Nevertheless, increasing light intensity consistently reduced the latency to calcium activation (**Fig. 4B**; 506.5 ± 67.6 s at 0.5 mW·cm^−2^, 244.2 ± 54.1 s at 1.0 mW·cm^−2^, and 87.6 ± 69.1 s at 1.5 mW·cm^−2^; *n* = 3). Together, these results show that delayed calcium activation under sustained sub-threshold physical stimulation is a general phenomenon and that its timing is systematically modulated by stimulus intensity.

**Fig. 4.**
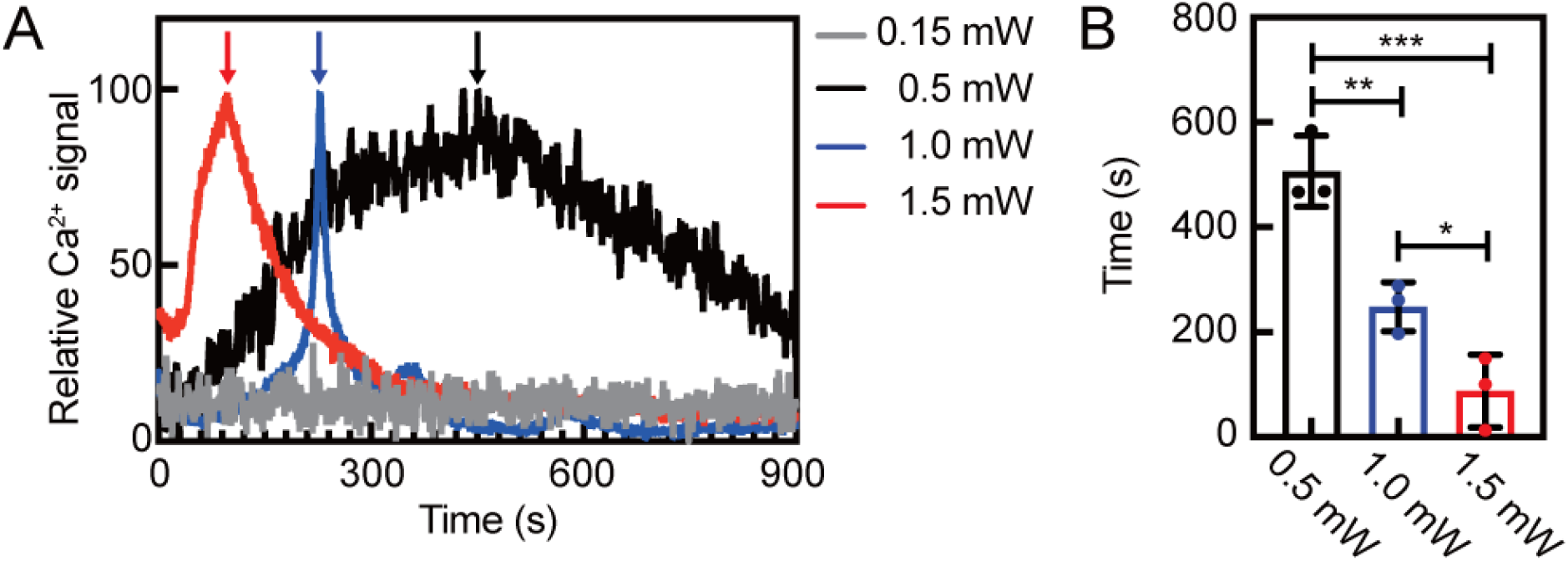
Intensity-dependent delay of calcium activation under sub-threshold light stimulation. (**A**) Representative fluorescence time traces of intracellular calcium responses in HUVECs subjected to sustained sub-threshold light stimulation. Traces are shown under three blue light intensities (0.15, 0.5, 1.0, and 1.5 mW·cm^−2^; 488 nm). Relative Ca^2+^ signals were calculated from background-subtracted fluorescence and subsequently min-max normalized, with values scaled 0 and 100, respectively. Calcium activation occurred with delayed onset under all conditions, with earlier onset at higher light intensities. (**B**) Quantification of latency to first detectable calcium activation under sub-threshold light stimulation. Higher light intensities were associated with reduced activation latency (0.5 mW·cm^−2^: 506.5 ± 67.6 s; 1.0 mW·cm^−2^: 244.2 ± 54.1 s; 1.5 mW·cm^−2^: 87.6 ± 69.1 s; mean ± s.d., *n* = 3). Latency was defined as the time required to reach peak calcium signal (indicated by arrows in panel a). Statistical significance was determined by one-way ANOVA followed by Tukey’s post hoc multiple-comparisons test to account for multiple hypothesis testing. Note: *n* = 3; \**p* < 0.1; \*\**p* < 0.01; \*\*\**p* < 0.001

## 3. Discussion

This study identifies temporally integrated, delayed cellular activation as a distinct response regime under sustained low-intensity physical stimulation. Under sub-threshold acoustic and light stimulation, endothelial cells did not exhibit rapid calcium activation but instead responded only after sustained exposure on the order of minutes under the tested conditions. The latency of this delayed activation was dependent on stimulus intensity, with stronger sub-threshold inputs reducing the time required to elicit calcium signaling. The observation of similar intensity-dependent behavior across distinct physical modalities indicates that delayed calcium activation reflects a general cellular response to prolonged weak stimulation rather than a modality-specific effect. These findings support a conceptual model in which cells integrate low-level physical inputs over time until an activation threshold is reached, resulting in delayed intracellular signaling (**Fig. 5**).

**Fig. 5.**
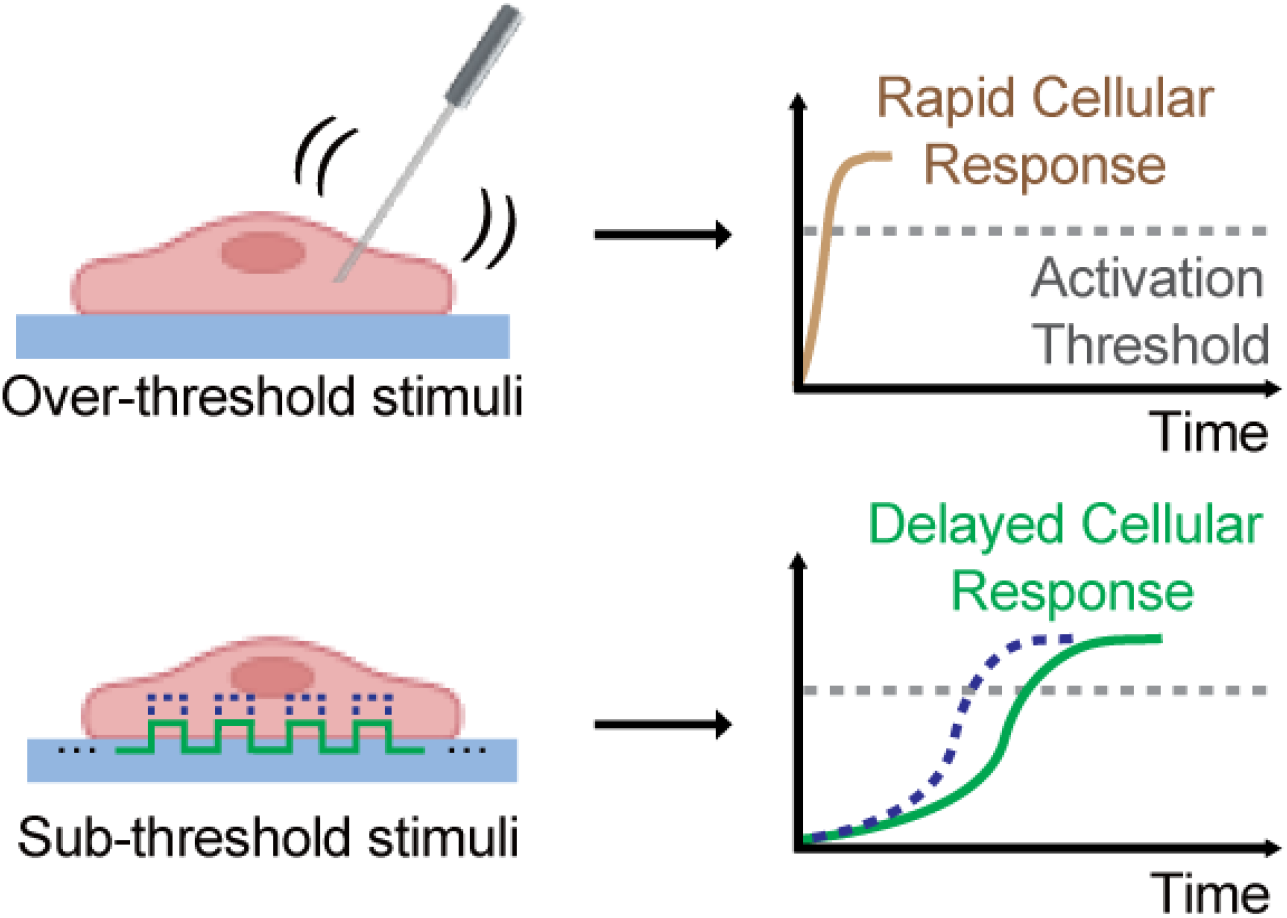
Proposed model of rapid and delayed cellular responses to over- and sub-threshold physical stimulation. Over-threshold stimuli (e.g., direct mechanical contact) exceed the activation threshold within a single event, producing a rapid intracellular response. In contrast, sustained, sub-threshold stimuli (e.g., acoustic or light stimulation) do not elicit immediate calcium responses are integrated over time, resulting in delayed intracellular activation once the threshold is crossed. Stronger sub-threshold inputs accelerate this integrative process, leading to earlier threshold crossing and reduced response latency. This model illustrates temporal integration as a cellular mechanism underlying delayed responses to sustained weak physical stimuli.

Over-threshold and sub-threshold physical stimulation elicit distinct calcium signaling regimes that differ in intercellular propagation and spatial organization. Direct mechanical perturbation exceeds the activation threshold within a single event and generates calcium signals of sufficient strength to engage cell–cell communication, resulting in coordinated, propagative responses. In contrast, sustained sub-threshold acoustic or light stimulation appears sufficient to activate intracellular calcium signaling at the level of individual cells but remains below the threshold required to affect neighboring cells. Under these conditions, calcium activation remains cell-autonomous and spatially dispersed, with responding cells emerging at variable and unpredictable locations across the population. This combination of weak intensity, delayed activation, and spatial unpredictability defines a signaling regime that differs fundamentally from that induced by strong, localized perturbations.

This signaling regime has particular relevance for weak environmental exposures such as chronic noise, which are applied broadly and persistently. In such contexts, cellular responses may instead arise sporadically at the level of individual cells through temporal integration of weak inputs. This property helps frame why long-term noise exposure has been associated with adverse health outcomes, even though clearly defined acute cellular responses have been difficult to identify. While the present study does not establish a direct connection between delayed calcium activation and downstream pathological pathways, calcium signaling is a central regulator of diverse physiological processes (*10, 11*). The discovery of temporally integrated, sub-threshold calcium activation therefore reveals an early intracellular response mode through which persistent environmental stimuli may influence biological systems, motivating further studies to determine how such dispersed signaling events propagate into longer-term biological effects.

Differences between light- and acoustically induced calcium responses demonstrate a broader challenge in identifying temporally integrated signaling under weak stimulation. Under sub-threshold light exposure, calcium responses exhibited broad temporal profiles and baseline trends that can complicate identification of clear activation events. Such features can obscure delayed responses in optical assays and have contributed to their attribution to nonspecific effects or experimental variability. In contrast, acoustically delivered stimulation, an orthogonal modality independent of optical excitation, produced comparable intensity-dependent delays. Together, these results demonstrate that delayed calcium activation reflects a genuine cellular response to sustained weak physical stimuli rather than an experimental artifact.

This study focuses on early intracellular calcium signaling responses in human endothelial cells, which are highly responsive to physical and mechanical cues and serve as key sensors of vascular and environmental stress. The experiments were conducted in a controlled in vitro setting and in a single cell type, providing a defined framework to isolate sub-threshold signaling behavior. Extending these observations to additional cell types and more complex biological contexts will be important to assess the generality and physiological significance of temporally integrated calcium responses to weak physical stimulation.

## 4. Experimental Section

### Design and fabrication of the stimulation chamber

An experimental chamber was constructed using polydimethylsiloxane (PDMS; Dow Corning, Midland, MI, USA) bonded to a glass coverslip (Thermo Fisher Scientific, Waltham, MA, USA) to define a cell-culture region adjacent to a transducer mounting region (Figure 1 and Figure S1). The PDMS prepolymer was prepared by mixing base and curing agent at a 10:1 weight ratio. To minimize light reflection from the PDMS structure during fluorescence imaging, carbon black powder (Thermo Fisher Scientific, Waltham, MA, USA) was incorporated into the PDMS prepolymer mixture at a final concentration of 1 wt% prior to curing (*21*). The mixture was poured into a custom 3D-printed mold, degassed under vacuum to remove trapped air bubbles, and cured at 100 °C for 1 hour. Each PDMS structure measured 4 × 3 × 0.5 cm (length × width × height). A circular cell-culture region (2 cm diameter) was positioned at a fixed distance of 0.2 cm from the transducer contact site to ensure consistent surface-wave propagation to the cell-culture region. The PDMS structure and glass coverslip were treated with oxygen plasma and irreversibly bonded to ensure leak-free sealing and mechanical stability. The glass coverslip served as the substrate for cell culture and surface-wave propagation.

#### Cell culture

Human umbilical vein endothelial cells (HUVECs; CRL-1730, ATCC, Manassas, VA, USA) were cultured in F-12K medium (Gibco, Waltham, MA, USA) supplemented with 10% fetal bovine serum (FBS; Gibco, Waltham, MA, USA), endothelial cell growth supplement (ECGS; Corning, NY, USA), and 0.1 mg/mL heparin (Sigma-Aldrich, Burlington, MA, USA). Cells were used at passage 5. The cell-culture region of the stimulation chamber was coated with 0.1 mg/mL of type I collagen (Cell Biologics, Chicago, IL, USA) and incubated at 37 ºC for 30 min, followed by three washes with phosphate-buffered saline (PBS). HUVECs were seeded at a density of 3 × 10^4^ cells/cm² and cultured to near confluence to form a continuous endothelial monolayer prior to physical stimulation.

#### Calcium indicator loading

Intracellular calcium dynamics were monitored using the fluorescent calcium indicator Calbryte 520AM (AAT Bioquest, Pleasanton, CA, USA). A 2 mM stock solution was prepared by dissolving 50 µg Calbryte 520AM in 20 µL dimethyl sulfoxide (DMSO; Invitrogen, Waltham, MA, USA) containing 2.1 µL Pluronic F-127 (Invitrogen, Waltham, MA, USA) to improve dye solubility. The stock solution was diluted in Hank’s Balanced Salt Solution (HBSS; Gibco, Waltham, MA, USA) lacking Ca^2+^ and Mg^2+^ to a final concentration of 4 µM Calbryte 520AM. The absence of extracellular Ca^2+^ during loading minimized background calcium signaling. All dye preparation steps were performed under light-protected conditions to prevent photodegradation.

For dye loading, culture medium was removed and 200 µL of the Calbryte 520AM working solution was added. Cells were incubated at 37 °C for 30 min to allow dye uptake, followed by addition of 1 mL of HBSS supplemented with 10% FBS and incubation at 37 °C for 45 min to permit intracellular de-esterification. Cells were washed three times with HBSS to remove residual extracellular dye and reduce background fluorescence. Fresh HBSS was added, and cells were equilibrated for 10 min prior to imaging to stabilize baseline fluorescence.

#### Fluorescence imaging and intensity analysis

Fluorescence imaging was performed using a Ti2-A inverted fluorescence microscope (Nikon, Tokyo, Japan) with 488 nm excitation and 528 nm emission detection. Images were acquired with an exposure time of 100 ms per frame, providing sufficient temporal resolution to capture rapid intracellular calcium changes. All calcium imaging experiments were conducted under consistent environmental conditions and identical microscope configuration (objective, exposure time, and acquisition rate) to ensure comparability across experiments. For no-stimulation controls and for mechanical and acoustic stimulation experiments, fluorescence imaging was performed using low-intensity blue light (488 nm, 0.15 mW·cm^−2^), which did not elicit detectable calcium activation. In light stimulation experiments, incident light intensity was varied (0.15, 0.5, 1.0, and 1.5 mW·cm^−2^) while all other imaging parameters were maintained constant.

Fluorescence intensity over time was quantified from small square areas centered on individual cells exhibiting the earliest detectable Ca^2+^ activation. Background fluorescence from a nearby cell-free area was subtracted to correct for background signal. For each cell, intensity values were normalized to a 0–100 scale using min–max normalization to facilitate latency comparison (Figures 3 and 4). For image presentation (Figures 2 and S3), contrast settings were applied consistently within each stimulation condition for visualization purposes. Quantitative analyses were performed on the original background-corrected intensity values.

#### Pulse-wave excitation and measurement configuration

To validate transmission of pulse-waves from the transducer to the cell-culture region, surface pulse-wave propagation along the coverslip substrate was experimentally characterized using a remote acoustic sensing setup. The cell-culture region of the stimulation chamber was filled with Hank’s Balanced Salt Solution (HBSS; Gibco, Waltham, MA, USA) to emulate biological incubation conditions. All measurements were conducted in the absence of cells.

Pulse-wave propagation was measured using a scanning laser Doppler vibrometer (SLDV; MSA-100-3D, Polytec GmbH, Waldbronn, Germany) equipped with a focused laser beam (∼4 µm spot size). To enhance optical signal return, retroreflective tape was attached beneath the center of the cell-culture region. The assembled chamber was supported on a vibration-isolating foam layer to minimize mechanical interference from the measurement stage (Figure 1).

Excitation was provided by an ultrasonic transducer (V133-RM, Evident Scientific, Tokyo, Japan) positioned at the transducer mounting region and aligned with the longitudinal centerline of the cell-culture region. A synchronized external signal generator supplied the driving pulse waveforms and ensured phase coherence between excitation and velocity detection.

#### Pulse-wave acquisition parameters and propagation analysis

Pulse-wave excitation was applied with a 70% duty cycle at pulse repetition frequencies of 220, 100, 50, and 10 kHz and at driving voltages ranging from 10 to 90 V to assess surface pulse-wave propagation across a range of excitation frequencies and amplitudes. Voltages up to 90 V were used for validation, while higher voltages were applied in cell experiment. Velocity measurements were acquired at a sampling frequency of 6.25 MHz, corresponding to a temporal resolution of 160 ns. For each pulse repetition frequency, the acquisition window was selected to capture at least ten complete waveform cycles. Each measurement point was averaged over 20 acquisitions to improve the signal-to-noise ratio. Spatial scanning was performed along a linear array of points spaced 480 µm apart, aligned with the longitudinal axis of the PDMS–glass substrate. Pulse-wave propagation toward the cell-culture region was evaluated by quantifying relative temporal phase lags between sequential scan positions using SLDV. A propagation toward the cell-culture region was confirmed by progressive delay in waveform arrival time between adjacent positions (Figure S1).

#### Statistical Analysis

Latency of intracellular calcium activation was quantified from fluorescence time-course data. For each stimulation modality, differences among stimulus intensities were analyzed using one-way ANOVA followed by Tukey’s post hoc multiple-comparisons test. Analyses were performed under the assumption of equal variance. All experiments were conducted with *n* = 3 independent biological replicates per group. Statistical analyses were carried out using GraphPad Prism 9 (GraphPad Software, Boston, MA, USA). Significance levels are indicated as follows: \**p* < 0.05; \*\**p* < 0.01; \*\*\**p* < 0.001; \*\*\*\**p* < 0.0001; ns, not significant (*p* > 0.05).

## Supporting information

Supporting Information_Soung-Biosignaling

## Acknowledgements

This work was supported by the Science and Technology Center New Frontiers of Sound (NewFoS) through U.S. National Science Foundation (NSF) cooperative agreement #2242925.

## Author Contributions

K. R., P.A.D. and M.K. conceived the project; Y.H.N., K. R., X. D., P.A.D. and M.K. designed experiments; Y.H.N., I.H, and M.K. conducted experiments; Y.H.N., I. H., K. R., X. D., P.A.D. and M.K. performed data analysis; Y.H.N. and M.K. wrote the original draft; All authors edited the manuscript.

