## Supporting Information_Soung-Biosignaling for "Temporal Integration of Weak Acoustic Stress Leads to Delayed Cellular Responses"

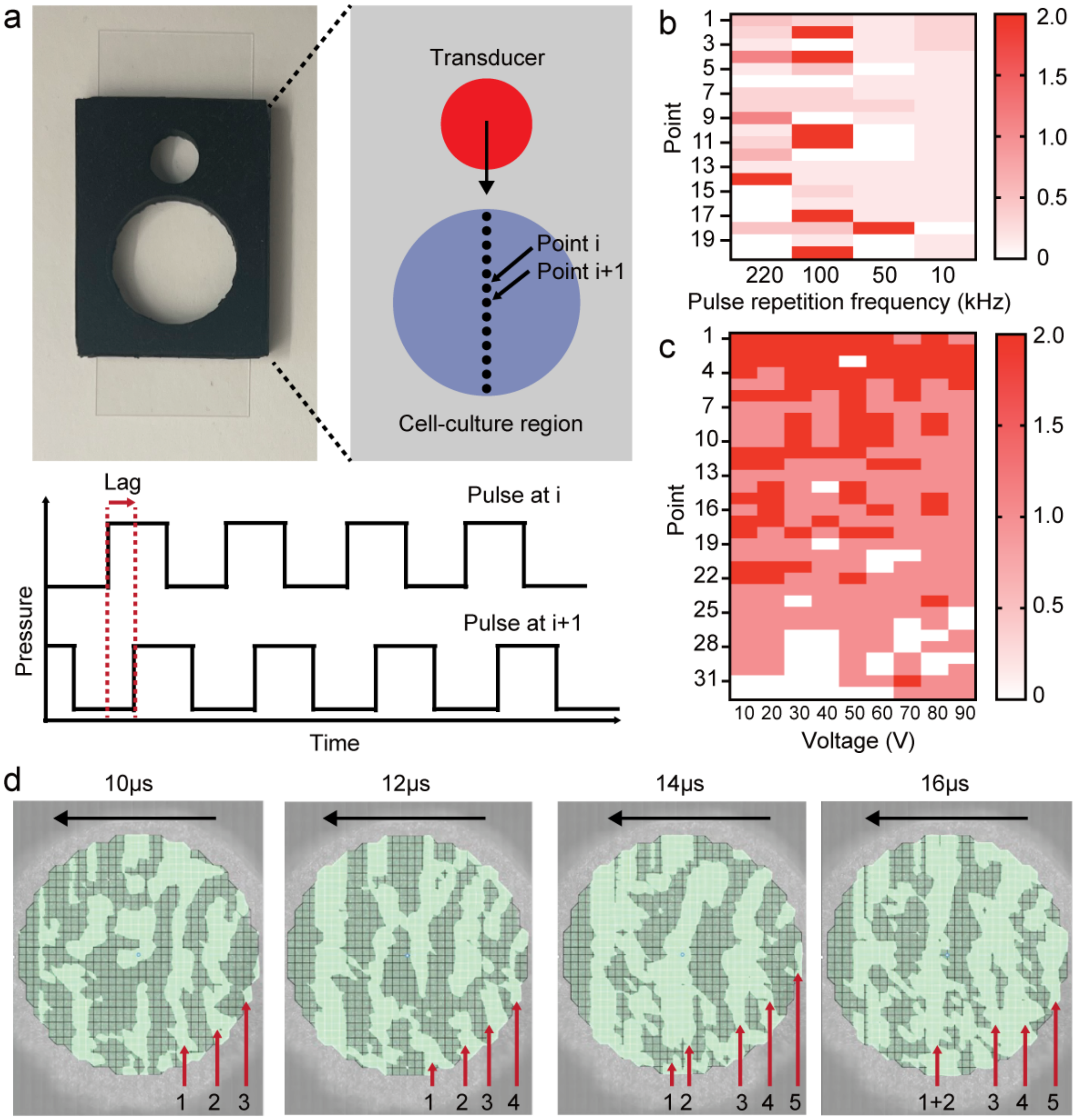


**Figure S1. Characterization of surface pulse-wave propagation from the transducer to the cell-culture region.** (a) Photograph of the experimental platform and schematic of the lag analysis used to evaluate surface pulse-wave propagation. Pulse waveforms were recorded at sequential positions along the substrate surface (Point i and Point i+1). The time difference between corresponding waveforms at adjacent positions was determined by shifting one waveform in time until the best alignment between the two signals was obtained (defined here as the temporal lag). When the waveform at the downstream position occurred slightly later than that at the upstream position, this delay (positive lag) indicated that the pulse-wave traveled across the surface toward the cell-culture region. When no systematic delay (zero or inconsistent lag) was observed, the coherent surface propagation pattern was not evident. (b) Heat maps of calculated temporal lag values under pulse-wave stimulation at different pulse repetition frequencies. Color intensity reflects the measured temporal lag between adjacent scan positions. Non-zero lag values indicate detectable surface pulse-wave propagation, whereas near-zero values indicate minimal or no propagation. Among the tested pulse repetition frequencies, 10 kHz exhibited the most spatially consistent propagation profile. (c) Heat maps of calculated temporal lag values under pulse-wave stimulation at different input voltages (pulse repetition frequency = 10 kHz). Across the tested voltage range, positive temporal lags were observed between sequential scan positions, confirming effective surface pulse-wave propagation from the transducer toward the cell-culture region. (d) Representative time-resolved laser Doppler vibrometry (LDV) images showing surface pulse-wave propagation within the cell-culture region under defined stimulation conditions (pulse repetition frequency = 10 kHz, input voltage = 80 V, 70% duty cycle). Measurements were performed in the absence of cells, while maintaining the same chamber geometry, buffer conditions, and excitation configuration used in cell experiments, to validate surface pulse-wave delivery to the culture region. The black arrow indicates the direction of propagation from the transducer, and red arrows denote sequential wavefront positions at increasing time points, illustrating temporal progression of the pulse-wave.

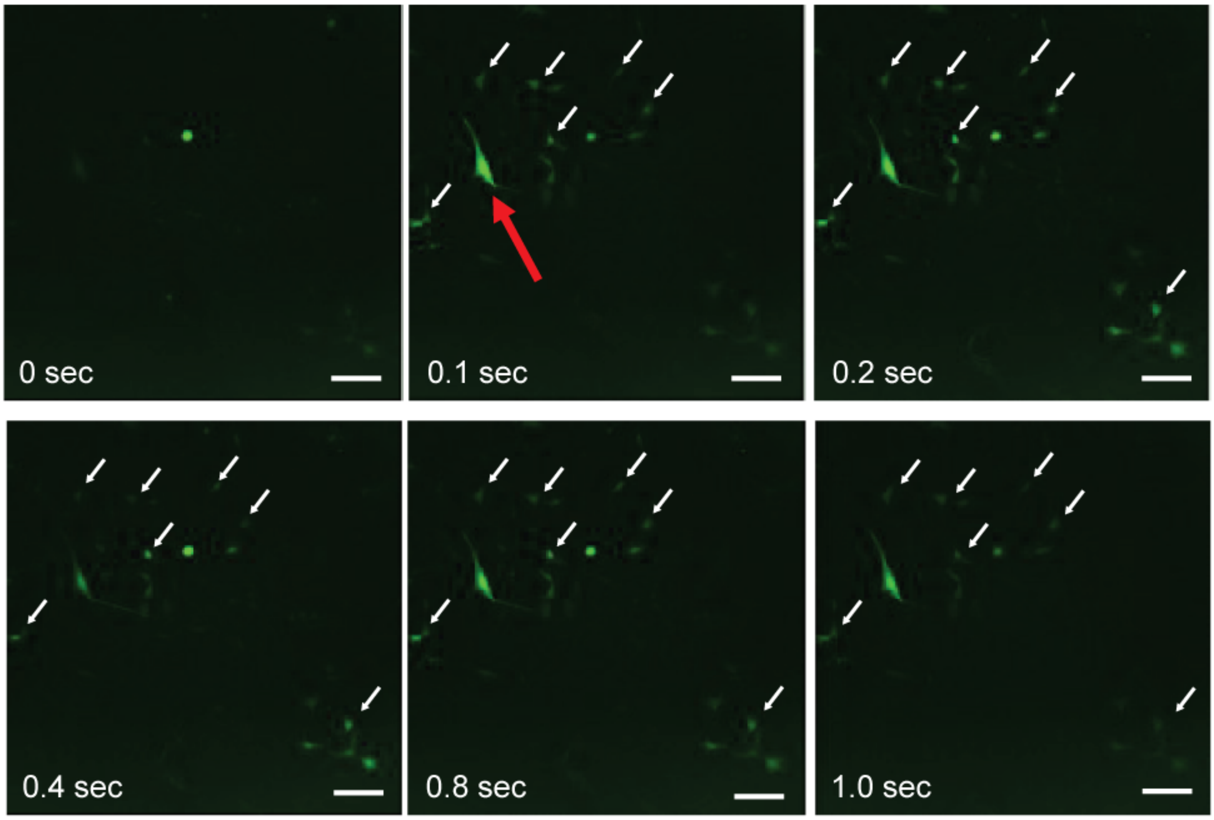


**Figure S2. Rapid intracellular calcium activation and intercellular propagation following direct mechanical stimulation**. Representative fluorescence images showing a microneedle contacting a single endothelial cell (red arrow). A rapid increase in intracellular calcium signal was observed in the contacted cell within 0.1 s following stimulation onset. Subsequent frames (0.2–1.0 s) show calcium activation spreading from the stimulated cell to neighboring cells (white arrows), consistent with intercellular propagation of the calcium signal. Scale bar, 100 µm.

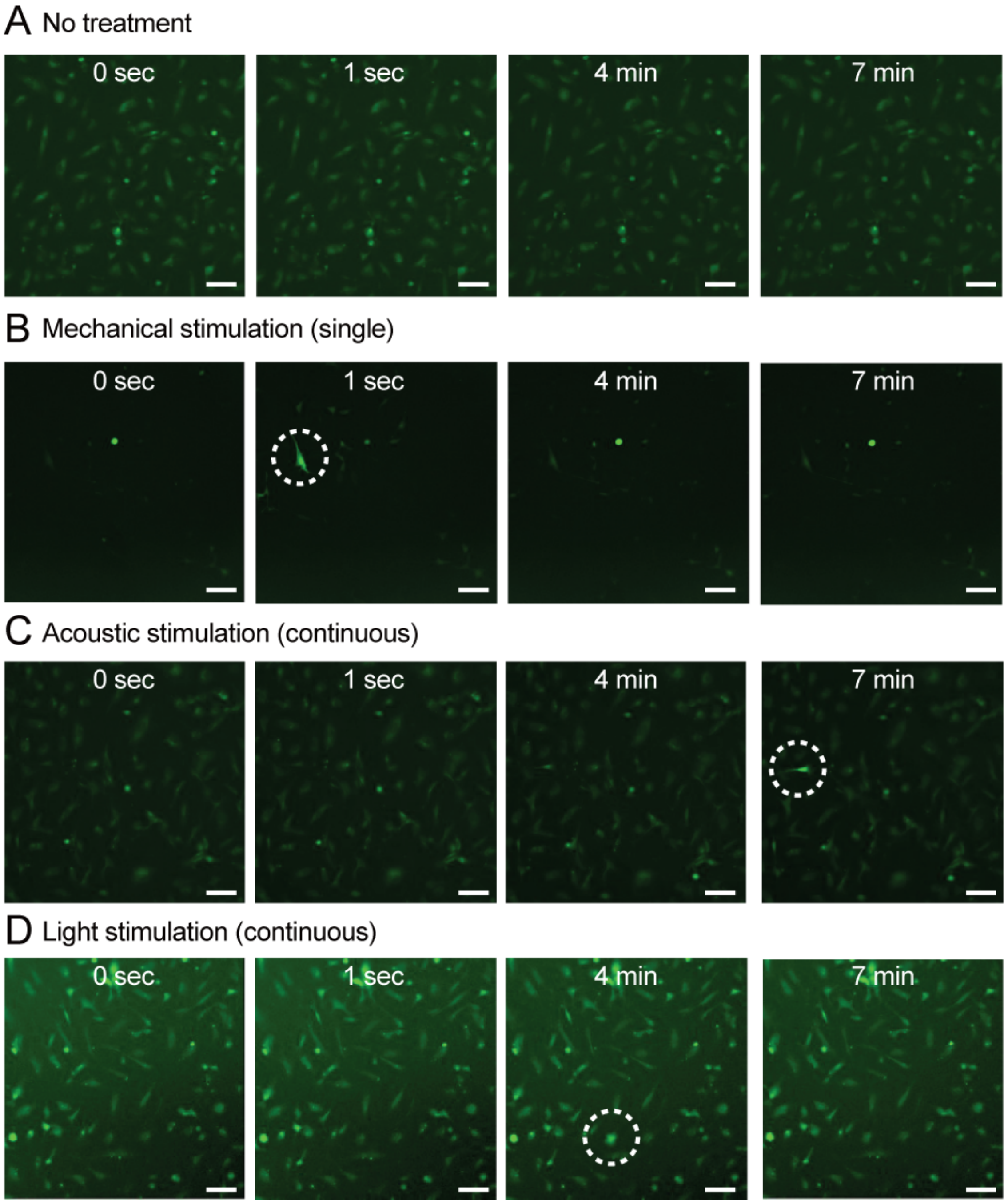


**Figure S3. Low-magnification overview of calcium dynamics under distinct stimulation conditions.** Representative time-lapse fluorescence images of Calbryte 520AM-loaded human umbilical vein endothelial cells (HUVECs). (a) Unstimulated control cells showing basal fluorescence over time. (b) Direct mechanical stimulation applied by a single microneedle contact, producing a rapid increase in intracellular calcium detectable within 1 s in the stimulated cell. (c) Continuous sub-threshold acoustic stimulation delivered as surface-propagating pulse-waves (10 kHz, 170 V input, 70% duty cycle), with intracellular calcium activation at later time points. (d) Continuous sub-threshold light stimulation (488 nm, 1.0 mW·cm$^{-2}$), with calcium activation at later time points. For (a)-(c), fluorescence imaging was performed using low-intensity blue light (488 nm, 0.15 $mW \cdot cm^{-2}$). As shown in (a), this imaging condition alone did not elicit detectable calcium activation. Image display contrast was adjusted uniformly within each stimulation condition for visualization clarity. Quantitative analyses were performed on background-corrected fluorescence intensity data as described in the Materials and Methods. White dashed circles indicate the locations where calcium activation first appeared. Scale bar, 100 μm.
